# MapCell: Learning a comparative cell type distance metric with Siamese neural nets with applications towards cell-types identification across experimental datasets

**DOI:** 10.1101/828699

**Authors:** Winston Koh, Shawn Hoon

## Abstract

Large collections of annotated single-cell RNA sequencing (scRNA-seq) experiments are being generated across different organs, conditions and organisms on different platforms. Transferring annotations from this growing database of single cell expression data to a new unannotated experimental dataset can accelerate insights into the underlying biology. There have been many approaches towards aligning and unifying such heterogeneous datasets. In our work, we recognized the need for a robust data driven distance metric to map annotation across datasets. Towards this aim, we applied a one-shot training approach, Siamese Neural Networks (SNN), to learn a distance metric that can differentiate between known annotated single cell types. Requiring only a small training set, we demonstrated that the SNN can perform predictions across different scRNA-seq platforms, identify novel cell types and transfer annotations across samples.

## INTRODUCTION

At a joint Cold Spring Harbor Laboratory/Wellcome Trust Genome Informatics conference in 2002, Sydney Brenner urged the development of a cell map to serve as “a framework to think of genomes and their products”^1^. This was further elaborated in 2010 where he defined CellMap as “a map of the molecules within cells and a map of the cells in the organism”^2^. The field on single cell analysis has evolved rapidly over the last few years primarily driven by the development of single cell RNA sequencing (scRNA-seq) which has led to community efforts like the Human Cell Atlas^3^ to enable a better appreciation of heterogeneity in complex tissues. This is paving the way for a better understanding of normal and pathological developmental programs. Many community tools have been developed that categorize heterogenous populations of cells, based on their gene expression, into types and states^4–6^.

Much of the effort conducted by these studies, involves careful clustering of cells, annotation using reference markers and subsequent validation of diverse cell states. This is often a manual and time-consuming process and the reliance on a clustering process can be subjective^6^. Furthermore, most deep learning techniques require large numbers of training examples to develop robust models. However, it is often not possible to obtain sufficient training examples to learn models for rare cell-types or disease cell states.

In this work, we developed MapCell, a deep learning method for classifying cell types at the single-cell level, with the aim to enable automated annotation of known cell types and discovery of new cell types. We demonstrated the ability to learn an independent cell type distance metric that can provide a quantitative measure of similarity when comparing pairs of single cells. Many machine learning algorithms depend critically on having a robust distance metric. However, standard distance measures such as Euclidean or cosine distance may miss subtleties in the properties of high-dimensional single-cell gene expression measurements.

We employed a form of one-shot learning, Siamese Neural Network (SNN), to learn a similarity function to differentiate between pairs of cells using their gene expression profiles as input. One-short learning is a classification task where one or very few examples of each class is used to train a model to make predictions of many unknown examples. Siamese Neural Network (SNN) is a popular architecture that has been developed for this task and it benefits from joint learning of both a feature representation space and a distance metric requiring few training examples to generate robust models. Siamese networks have already been used in areas like signature verification^7^, image recognition^8^ and facial recognition^9^, where the number of training examples for each individual class is limited and the number of classes is dynamically changing. This makes data collection and retraining costly. We find an analogous challenge in distinguishing cell types and states which can exist along a continuum and finding sufficient training examples for each state is difficult for standard deep learning techniques.

To demonstrate the use of SNN for single cell analysis, we focused on a comprehensively labelled dataset derived which catalogued single cell data of myeloid cells originating from matched peripheral blood and tumors of 7 non-small-cell lung cancer (NSCLC) patients^10^. Training of the SNN required only 30 training examples per cell type.

The process of training, deployment and validation of SNN distance metric on single cell RNA-seq (scRNA-seq) expression data generates a reduced dimension embedding space that we used to visualize the similarities between cells. In addition, by using SNN distance metric in comparing cell pairs, cells from novel types which are not represented in the training data result in constantly large distances when compared across all reference cell types, a signature which can be used for novelty detection. We also showed that the learned distance metric is generalizable such that cell types unseen in the training set can still be distinguished from other cell types. This allows for refinement of the model simply by adding new cell-types to the list of reference cell types without requiring re-training of the model.

We show that a unique embedding space for each patient in the dataset could be derived due to the ease of training new models using SNN. The personalized embedding space is a form of a digital twin that captures the personalized information of cell types within each sample or person. We contrasted these patient models by deploying all models onto a single patient to illustrate the consistency of these models in annotating common cell types and the differences that arise when particular cell types are missing from individual models. We also trained a model using the Human Cell Landscape dataset^11^ which consists of a wider survey of cell types to demonstrate the scalability of SNN. Lastly, we demonstrated the transfer of annotation between mouse and human data sets by mapping orthologous gene inputs across mouse and human data.

## RESULTS

### Siamese Neural Network layout for representation of single cell gene expression

The network architecture employed in this study is illustrated in Figure 1. Our Siamese neural network consists of two identical subnetworks with shared weights. This subnetwork consists of a 3-layer neural networks with 512, 512 and 32 nodes respectively. Dropout layers are introduced between layers to improve the generalizability of the embedding space.

**Figure 1.**
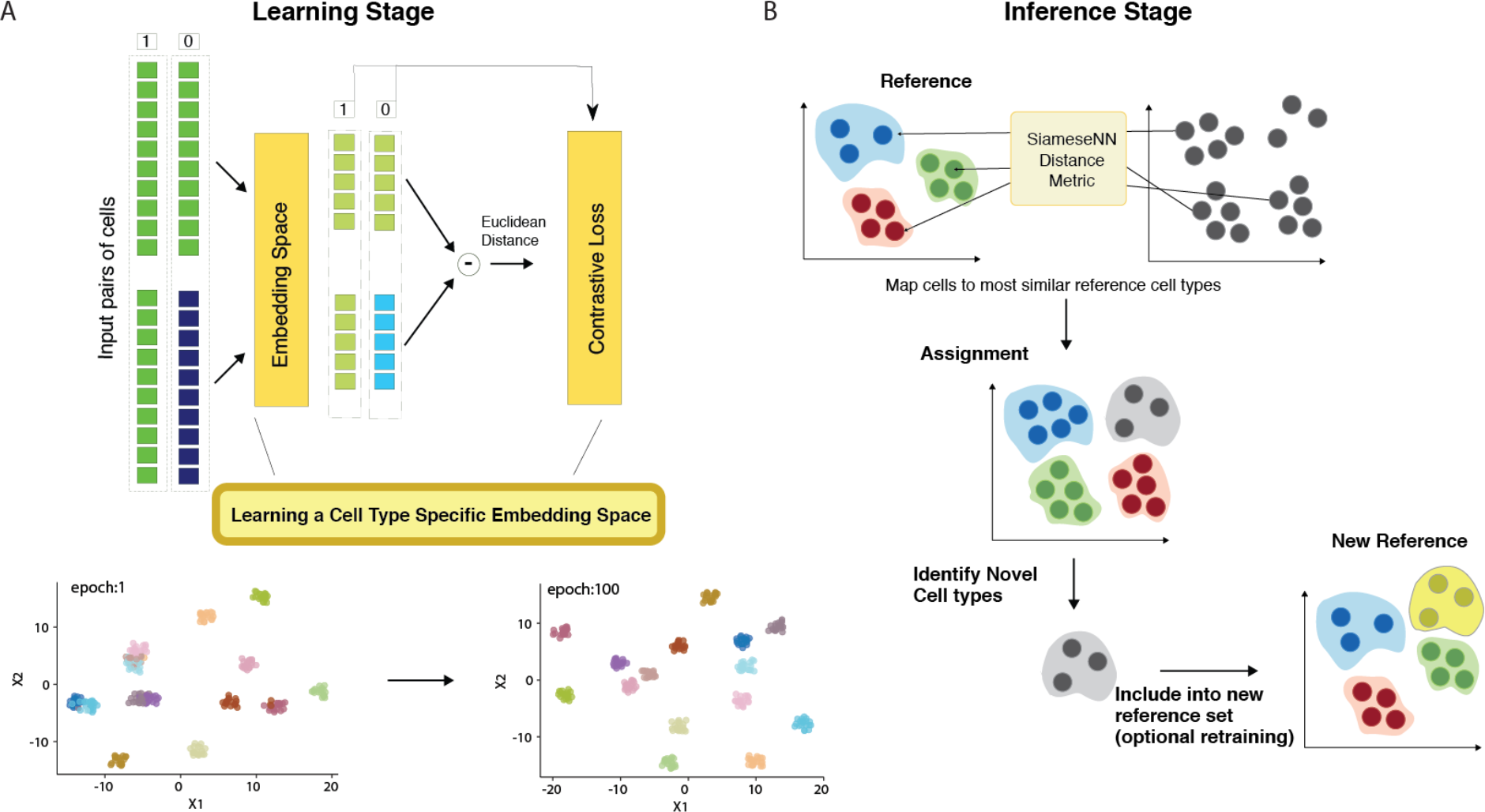
Overview layout and architecture of Siamese neural network. A. (Top) Siamese network architecture (Bottom) Low-dimensional representation of embedding space. B. The process of the Siamese Network inference. Each cell in the sample set is compared using the SiameseNN metric to a reference set of cells used in the learning stage. The assignment is made to the closest reference type. Cells that do not meet the threshold are flagged as novel cell types. These novel types can be reincorporated into the training set to generate a new SNN or included in the reference set without training.

To prepare the inputs for training, the counts of the most highly expressed gene is used to scale all other genes to ensure that input values are scaled between [0,1]. Generative functions were used during training to generate pairs of cells across cell types and each pair is fed into one of two identical subnetworks. Subsequent optimization was performed using a contrastive loss function. The training process can be visualized by examining the output of the last layer composed of 32 neurons using heatmap and UMAP dimension reduction visualization (Figure 2). The training set contains the same cell types originating from different tissues: peripheral blood and tumor. In the initial training epochs, cells from different cell types are already differentiated in the final neural net layer. Similar cell types found in different tissues were resolved as training further progressed. For example, B-cells from peripheral blood and tumor, were clustered together in epoch 1 but subsequently resolved in epoch 9 (Figure 2A). Similarly, in the embedding space, tumor NK and T cell were more similar to each other than their peripheral counterparts in epoch 9 but subsequently resolved by epoch 100 (Figure 2A). We can also observe the firing patterns of the neural network using a heatmap representation (Figure 2B). We see that firing patterns become more discrete as training progresses. The heatmap also reflects the complexity of the learned neural network. The significant number of unused nodes (blue squares in Figure 2B) suggest that a less complex network could be employed for further performance optimization.

**Figure 2.**
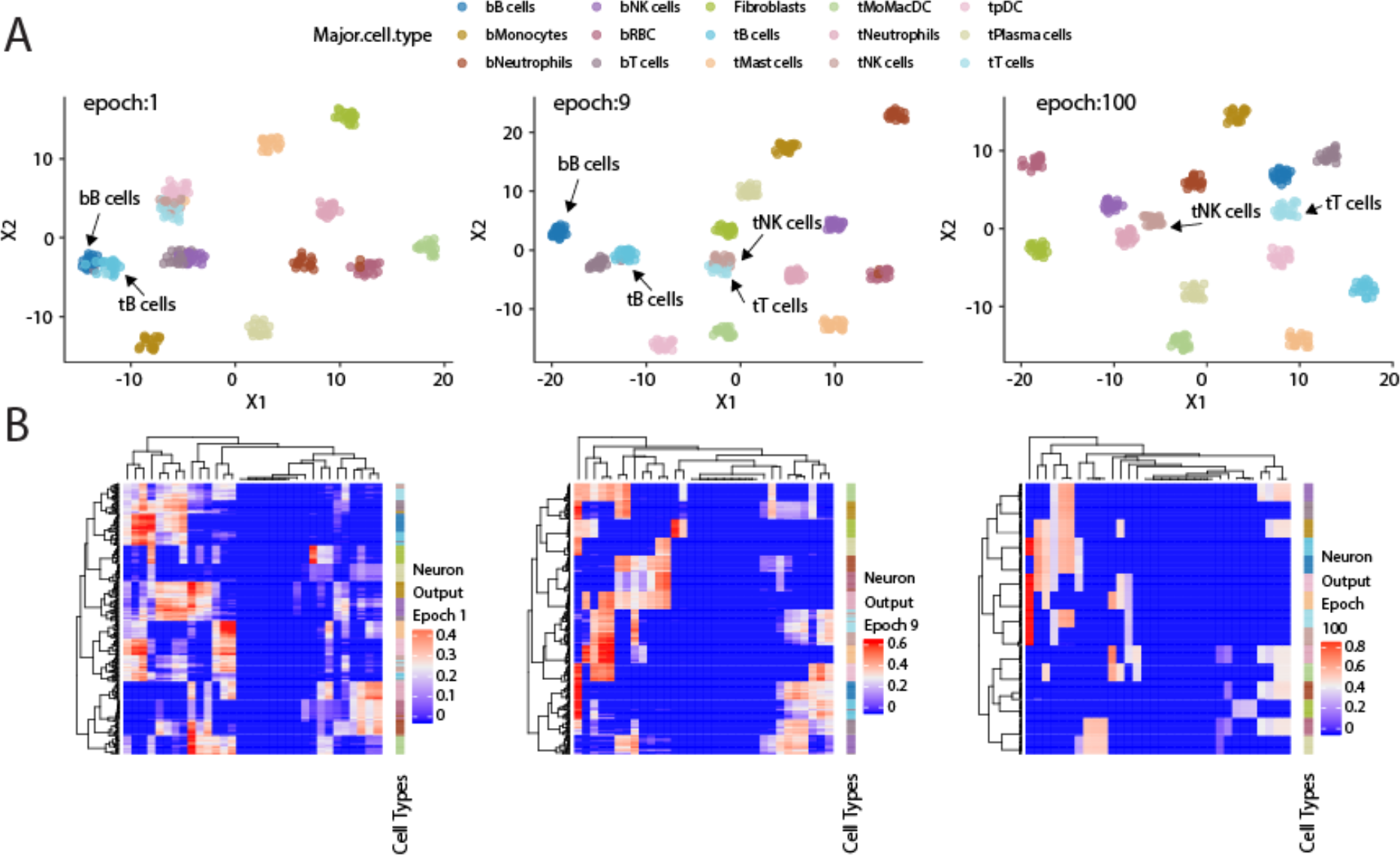
Visualization of training phase of Siamese Neural Network. (Top Row) Umap visualization of the embedding space projection in the last neuronal output at different training epochs. Example cell types that are better resolved, as measured by increased spatial separation in the embedding space over increasing training epochs, are indicated with arrows. (Bottom Row) Heatmap representation of the neural network firing patten where each row is a cell and and each column a single neuron in the final output layer.

### Using SNN for cell-type annotation

To illustrate the generalizability of using SNN distance, we proceeded to employ SNN distance trained on inDrop scRNA-SEQ platform^12^ to annotate a PBMC10K dataset generated by the 10× Chromium system, a different scRNA-seq droplet platform (10× Genomics). The 10× Chromium dataset provide by 10× Genomics included simultaneous cell surface protein measurements using oligonucleotide-tagged antibodies that provide an orthogonal validation of cell identity.

The annotation process works by comparing each cell in the PBMC10K evaluation set to 20 reference cells used in the training set. The cell type with the closest average distance is assigned (Figure 3A). Five major cell-types labels (bT cells, bB cells, bMonocytes, bNK and tPlasma cells) from the reference dataset are mapped onto the evaluation data (Figure 3B). The corresponding annotated cells clusters exhibited the canonical cell surface makers as illustrated by overlaying protein expression levels onto cells in the RNA defined t-SNE space (Figure 3C). The protein boundaries between cell clusters agree with the cell type boundaries annotated by the SNN. Notably, for T Cells, B Cells, Monocytes and NK cells, the PBMC10K cells were mapped to the corresponding blood-derived cell types rather than the tumor-derived cell types. For plasma cells, the reference only contained tumor-derived plasma cell types and were mapped accordingly.

**Figure 3.**
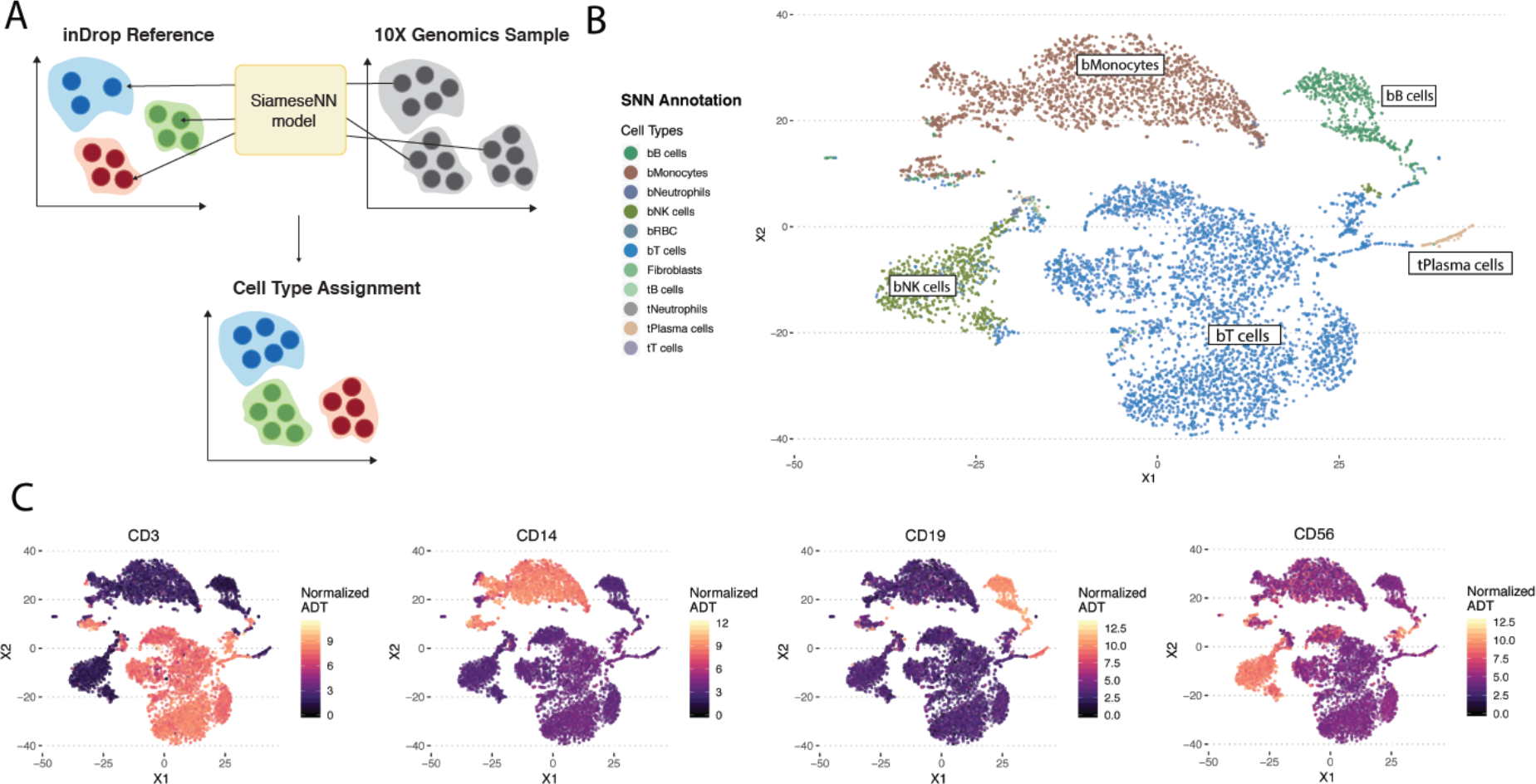
Workflow of the annotation of cell types using SNN. A. Each cell to be annotated (10× Genomics sample) is compared to 20 cells from each reference cell type using the learned SNN model. The reference cell type with the closest average distance is assigned to the cell. B. t-SNE plot of SNN annotated scRNA-seq data from 10× genomics. **C.** Cell surface protein expression mapped onto the respective cells in the same t-SNE space. Protein markers exhibit similar boundaries as the predicted annotations by SNN model.

### SNN distance is a better contrastive distance metric than cosine and Euclidean distances

We contrasted the SNN distance metric against commonly used Euclidean and cosine distance metrics. Twenty cells from each independently annotated cell type were compared pairwise against twenty cells across other cell types. SNN distance is able to resolve similar cell types better than Euclidean or cosine distance. SNN distance was small and showed a non-linear drop-off for similar cell pairs. This drop off is consistently observed for SNN distance metric across cell types. We quantified this using a signal-to-noise statistic (Figure 4D). The gain in signal is most pronounced when comparing red blood cells (RBC) across all other cell types (Figure 4B, 4D). As RBCs are biologically distinct from the other white blood cell types, we see a much smaller distance for RBC-RBC comparisons in contrast to other cell types. This is also reflected in the lower signal-to-noise ratio for similar cell types. This is especially evident for T and NK cells. While cosine and Euclidean distances were unable to unambiguously distinguish between T and NK cells types, SNN defined a clear demarcation between the two cell types while still ranking them as the two closest cell types. This feature of SNN was also useful for distinguishing different cell states. It is known that lymphocytes that infiltrate the tumor have a distinct cell state from lymphocytes found in peripheral blood^13^. With both Euclidean and Cosine distance, tumor B-cells (tB cells) were not well-distinguished from peripheral B cells (bB cells) while SNN distance clearly distinguished tB cells from bB cells (Figure 5). Taken together, we have shown that SNN distance is a robust metric for both cell type and cell state comparisons.

**Figure 4.**
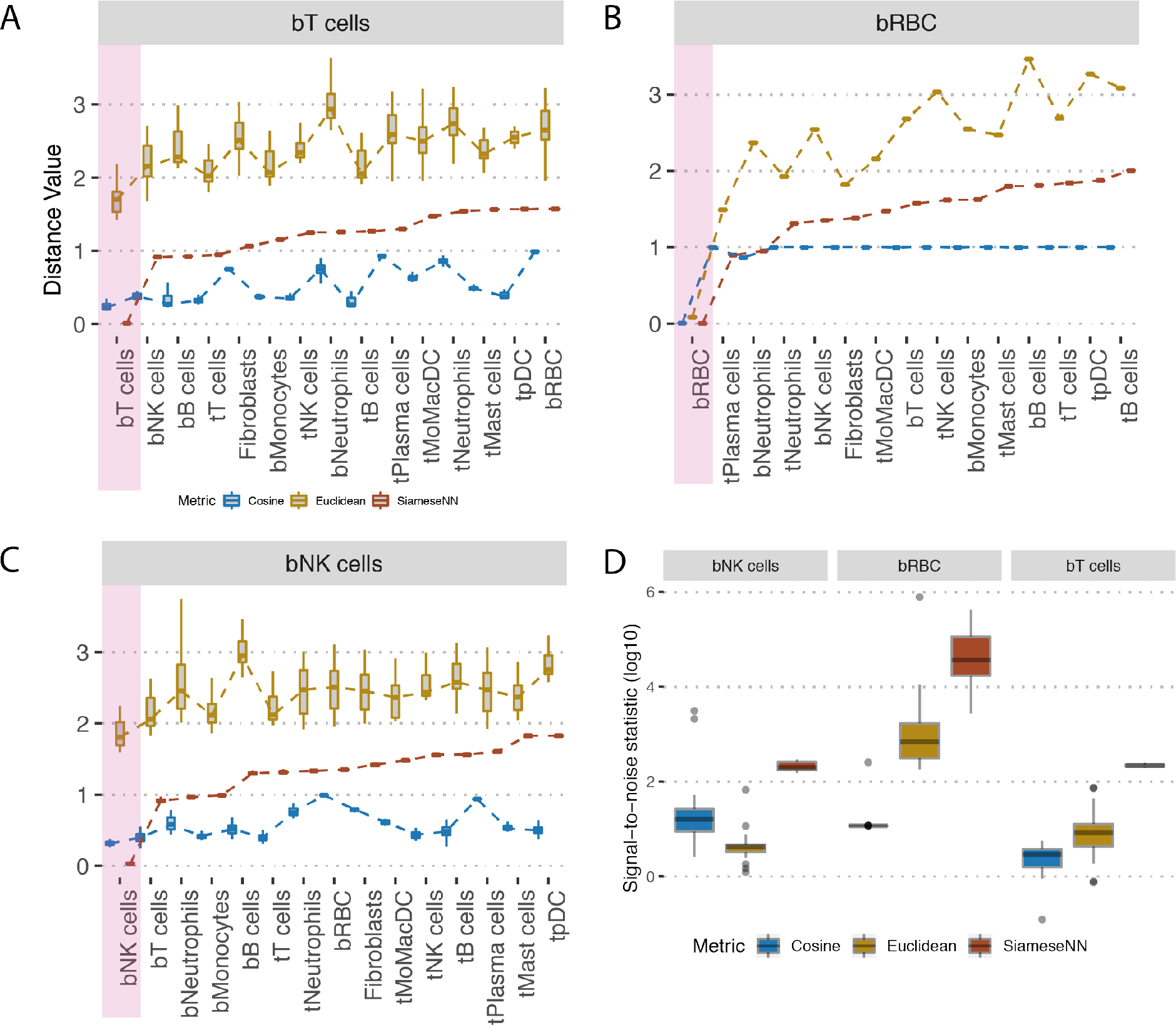
Contrasting Cosine and Euclidean against SNN distance for T-cells, RBC and NK cells. A-C. Boxplots showing distribution of pairwise distance scores between each annotated cell type from the test set against annotated cell type from the training set. The x-axis is ordered from left to right by the SNN distance values in ascending order. Colored red box indicates the reference cell type with the average closest distance. D. Boxplot showing the distibution of signal-to-noise values for each predicted cell type.

**Figure 5.**
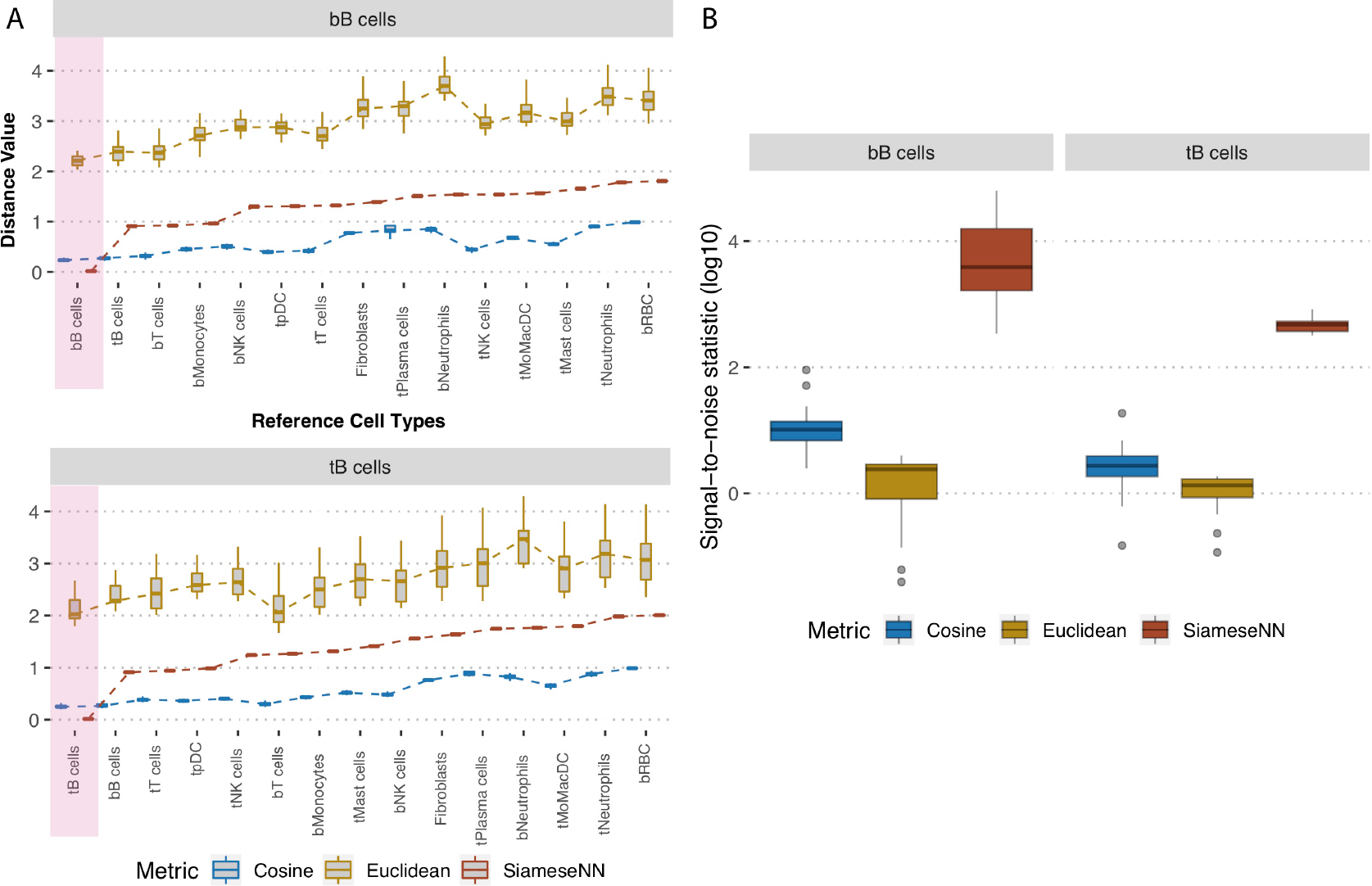
Contrasting Cosine and Euclidean against SNN distances for distinguishing cell state. A. Peripheral B cells (bB cells) and tumour derived B cells (tB cells) from the test set are compared against the reference cell types in the training set. B. Boxplot showing the distibution of signal-to-noise values for the different distant metrics for blood and tumor derived B-cells.

### Identification of novel cell types

As larger surveys of single-cell experiments are performed, we need to account for cell types and states that are not present in the training data set. For the purpose of refining the annotations and SNN model, it is more desirable to flag these novel cells rather than assign them to the closest cell type found in the training set. We examined whether the SNN distance metric can be used in novelty detection. We selected cells from a patient with three cell types (Type II cells, endothelial cells and patient-4 specific cells) that were not present in the training set and compared them against the reference cell types in the training data (Figure 6A and 6B). Predicted cell types were largely in agreement with human annotations (Figure 6C). We defined a novelty filter that will flag a cell as novel when the minimum distance computed across all cells is less than two standard deviation from the rest of comparisons. We found five regions that contained a high number of novel cell types. Expectantly, three of the five regions contained cell types not seen in the training set (black boundary, Figure 6D). The other two regions were found in the MoMacDC and T-cell cluster (green boundary, Figure 6D). Upon closer examination, we found that the MoMacDC cluster was comprised of clusters of subtypes ^10^ that were under-represented in the training set. As a result, the network did not recognize these cells as belonging to the MoMacDC cluster. We trained a new network that used the subtype labels to generate additional pairs of cells from these subtypes for training. This resulted in a more comprehensive training set and a better classification result (Figure 6E). The MoMacDC and T-cell clusters were no longer flagged as novel while the unseen training examples remained flagged as novel (Figure 6F). We used an alluvial plot to visualise the change in mapping of cell annotations after subtype training (Figure 7A). In agreement with the UMAP visualization, we see that after training on the subtype labels generated training set, we find a better mapping of the tMoMacDCs and tT cells (Figure 7B). This demonstrates a process where a novel cell type can be automatically flagged by the SNN for human inspection. This cell type can then be incorporated into the reference database for futher training. We did also observe however, that a minority of the tTcells which were classified correctly before, were misannotated to a different cell type. This could reflect the quality of the underlying published sub cell type annotation or insufficient sampling of training examples from the subtypes that led to overfitting of the model.

**Figure 6.**
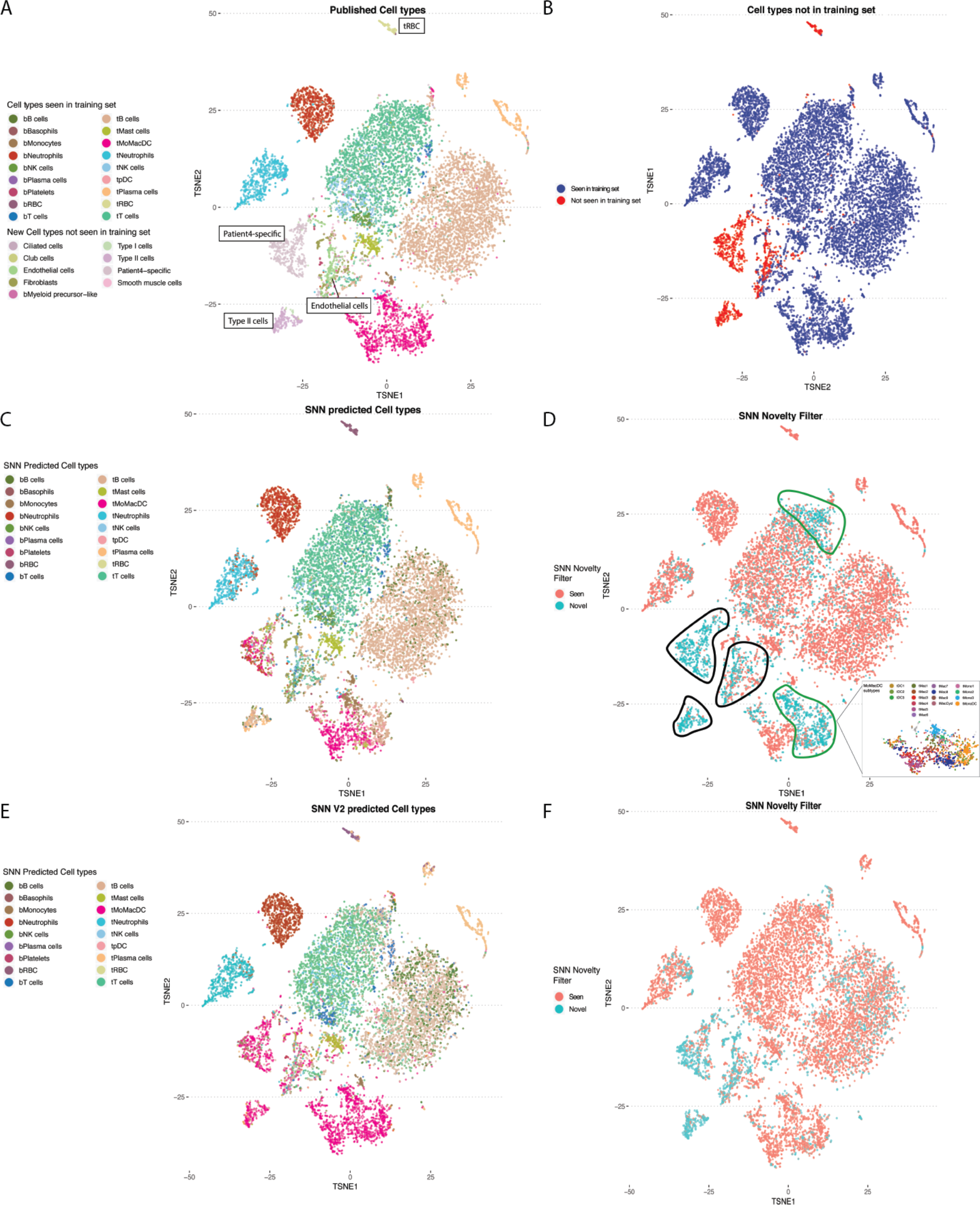
Siamese network distances for novelty detection. **A.** TSNE plot of single-cell RNA-seq NSCLC sample color by cell types annotated in study [3]. Cell types not present in the training data set are labled in boxes (tRBC, patient-4 specific cells, endothelial cells, Type II cells) **B.** TSNE plot of the same data. Highlighted in red and blue are the cells which were classified as cell types found and not found in the training set respectively. **C.** TSNE plot colored by SNN predicted cell types. **D.** TSNE plot colored by SNN novelty filter. Cells marked by black lines are flagged novel cells not seen in training set. Cells marked by green lines are flagged novel cells seen in training set. Callout box are the MoMacDC cluster colored by annotated subtypes **E.** TSNE plot colored by cell-type classification after re-training to include different sub-celltype examples. **F.** TSNE plot color by SNN novelty filter after re-training with sub-celltype examples.

**Figure 7.**
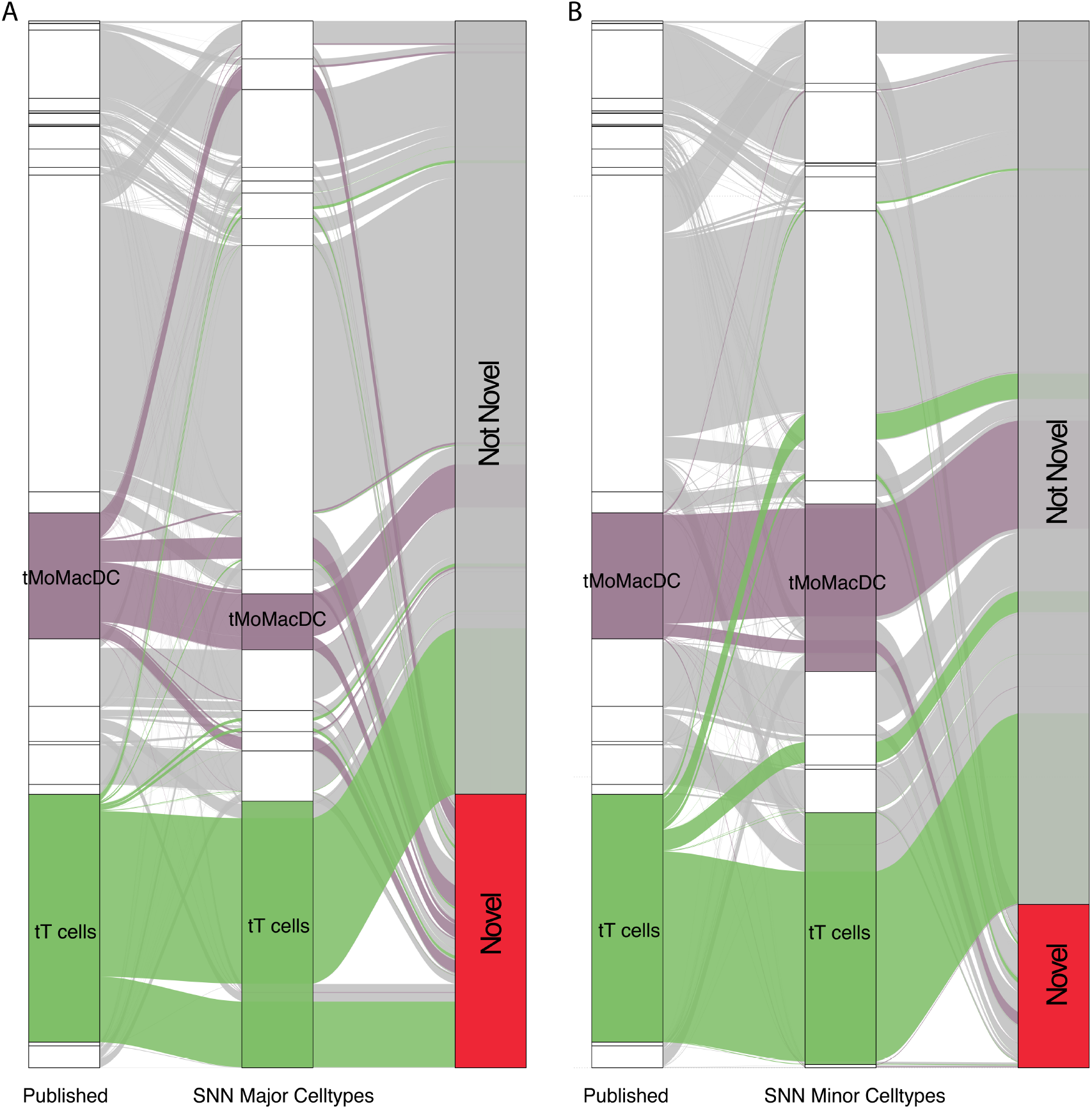
Alluvial plot depicting the switch in novelty status and annotation status when incorporating left out subtypes during training of SNN models. A. Mapping of cell types based on SNN trained on major cell type selected training examples. B. Mapping of cell types based on SNN trained on minor cell type selected training examples. Addition of omitted minor clusters of cell types redirects the annotations from novel to identifiable, and each to its respective expected human annotated states. The process depicts the capability of the SNN network to be used as an novelty detector as well as the plasticity of such a process to allow for subsequent update of novel classes.

### Siamese derived embedding space retains capacity to distinguish untrained cell types

Requiring a retraining process is a computationally intensive process. We explored whether the contrastive nature of Siamese network learns a general function that can be applied to new cell types without re-training. The intuition is that if sufficient diversity of gene expression measurements across different cell types are seen, the network would learn to weight different sets of genes representing pathways. These would enable new cell types, which have different combinations of pathways expressed, to be compared. Since there are unique cell types to particular patient groups in the Zilionis study^10^, we trained on one patient set (Fig. 8A) and projected cell types that the network was not trained on into the SNN-derived embedded space (Fig. 8B). We observed that the learned embedded space retains a general capacity to distinguish previously unseen cell types during training into separate clusters. To further validate the generalizability of the feature vectors in this embedding space, we generated a K nearest neighbor graph network using 20 cells from the trained cell types. We added to this graph the novel cell types that were not previously used for training and showed that distinct new cell types formed new clusters (Type II cells, Endothelial cells, Fibroblasts) while enucleated red blood cells from tissue or peripheral fraction were indistinguishable (Fig. 8C).

**Figure 8.**
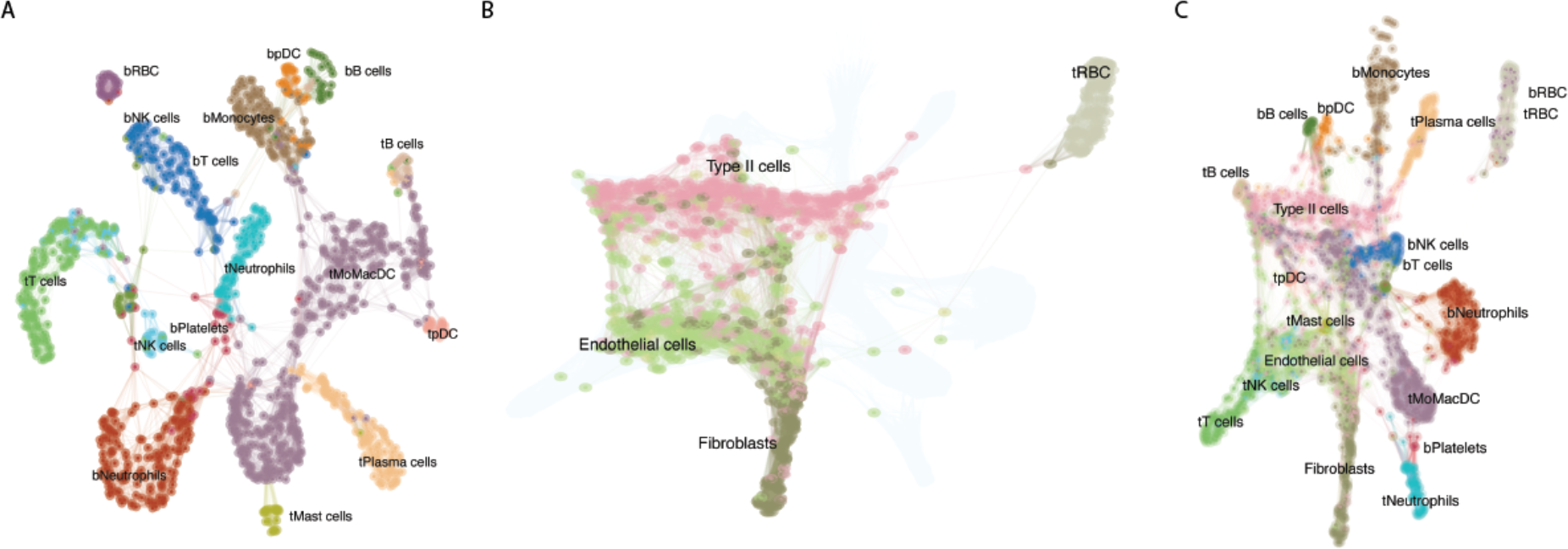
Siamese derived embedding space. **A.** K-nearest neighbor graph network of Siamese Network embedding space trained on a single patient. **B.** Projection of cell types not trained in the initial network onto embedding space. **C.** K-nearest neighbor graph network of Siamese Network embedding space with new cell types incorporated.

### Digital Twins and the scalability of SNN to accommodate large cell types numbers

Each patient profiled in the lung cancer dataset^10^ contained different numbers and diversity of cell types. To test the robustness of SNN models, we trained a unique model for each patient, leaving one patient out for validation. We treated each individually trained model as a pseudo-digital twin of the original patient. An alluvial plot is used to visualize the consistency and differences in annotations using personalized embedding spaces (Figure 9). We compared these personalized models against a model developed with a generalized embedding space that is capable of contrasting a large diversity of cell types. We obtained data from the Human Cell Landscape (HCA)^11^ that comprises a wide survey of cell types derived from about 50 different tissues. There are close to 700,000 cells in the data with 384 cell types and we sampled cells from cell types that are represented by at least 30 cells. The sampled cells were similarly used to generate pairs of contrasting cells in the atlas for training the Siamese network. We used this network to annotate the held-out sample. This demonstrated the scalability of the SNN architecture and its capability in accommodating cell types numbers on the order of an entire human cell atlas. Concordance of the major cell types were observed when comparing the annotations from the patient’s model as well as the HCL model. We also observed that the HCL model did not distinguish between T-cells of blood and tumor origin likely because these contrasting cell types were absent in the HCL dataset.

**Figure 9.**
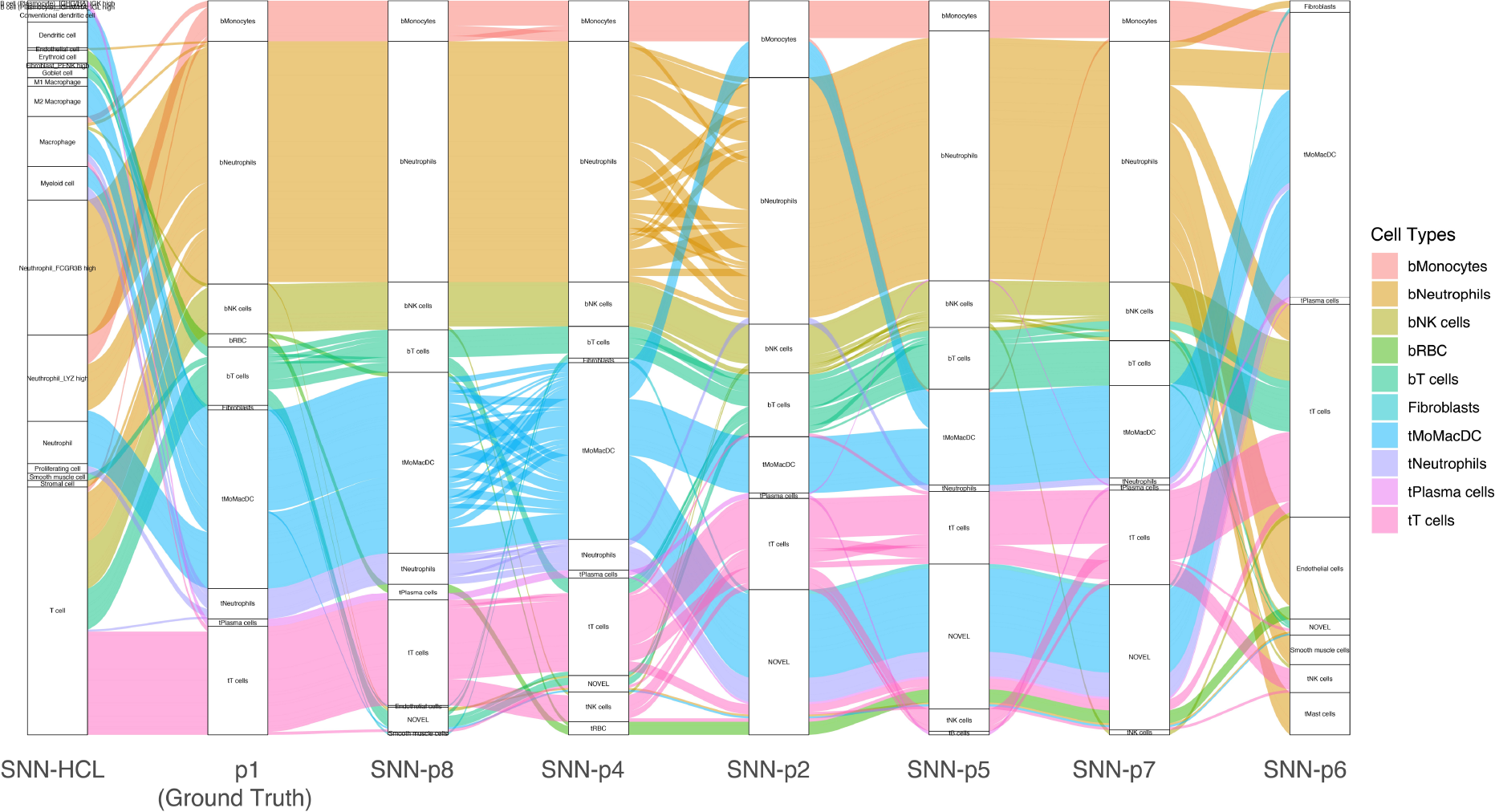
Alluvial plot of Patient 1 Annotations via individually trained models derived from other patients and the Human cell Landscape. Each column reflects the annotation of patient 1 data by various trained models using the SNN architecture. The leftmost column reflects annotations by SNN-Human Cell Landscape model. The second column represents the original annotation for patient 1 [10]. The remaining columns represents annotations by models trained with different patient data. In the leftmost column, we see that the human cell landscape based annotations were able to annotate T-cells accurately however it was unable to distinguish tumour derived T-cells from peripheral blood T-cells compared to other patient-SNN models.

### Interspecies annotation transfer

While the availability of single-cell genomics makes it possible to profile cells form different organisms, it is still a costly endeavor to generate atlases for multiple non-model organisms. Next we tested whether the SNN can be used to transfer annotation to a related species. Mouse genes were mapped to human orthologs and SNN prediction was performed on single-cell RNA-seq data of PBMC from a healthy mouse using a human reference. We showed that we could successfully annotate the mouse sample using the SNN trained from human data (Fig 10).

**Figure 10.**
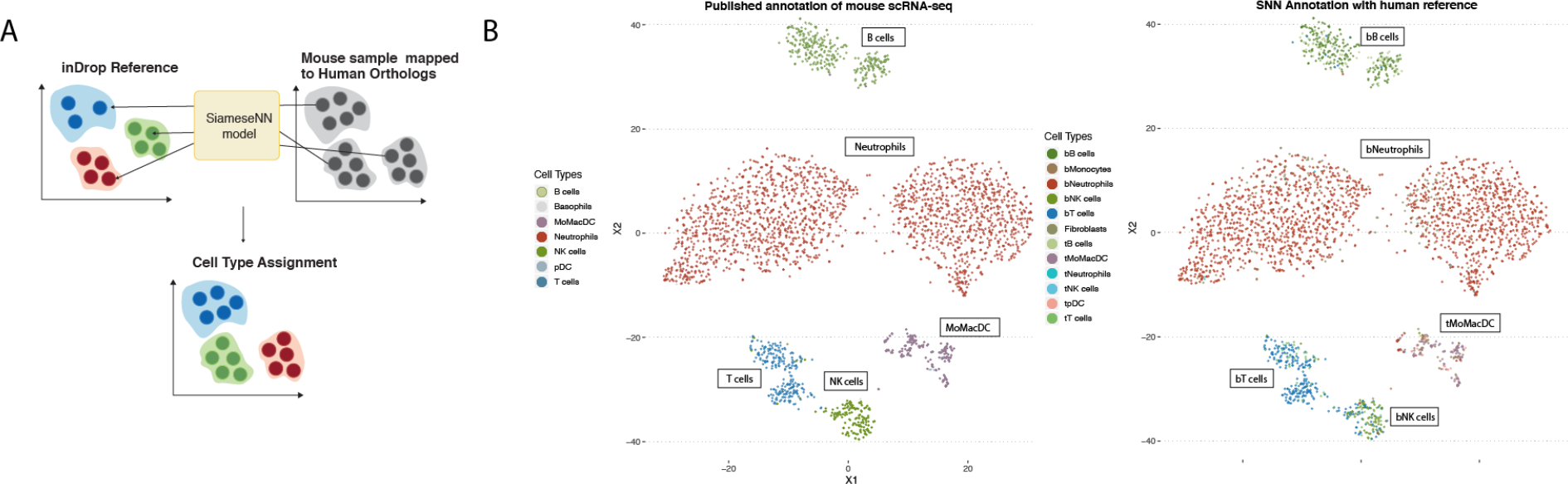
Inter-species annotation transfer. A. Transfer annotation learned on human to mouse. Mouse genes were mapped to human orthologs and SNN cell-type prediction was performed on mouse scRNA-seq data **B.** (Left) Published annotation of mouse scRNA-seq data (Right) SNN predicted cell types using human reference.

## DISCUSSIONS

We demonstrated the application of Siamese networks as a similarity function and demonstrated its usage in annotating cell types from single-cell RNASeq experiments. Training with this neural architecture requires only a small number of representative cells (30 in this study), making it ideal for learning of cell features of potentially rare cell types or transient states. Despite the small training set, we demonstrated that the SNN can perform predictions across different scRNA-seq platforms, identify novel cell types and transfer annotations across species. Our SNN-derived distance metric is robust compared to Euclidean and Cosine distance. We demonstrated that it can serve as a generalized metric for making comparisons for cell-types not seen in the training set. This allows the addition of cell-types to the SNN without the need for re-training. Furthermore, this distance metric can be integrated with other machine learning algorithms that employ distance metrics such as K-Nearest Neighbor (KNN) for rapid deployment. Future avenues of research include would include using similar a similar approach of trained models to compare samples across different timepoints. In our work, we have deployed models comparing different patients as a means of a novelty detector of cell types. It is conceivable that such an approach to be applied for a single patient comparing multiple timepoints against a baseline model. Such a baseline model can be thought of as a digital twin of the patient capturing the diversity of the patient cell types and states in the trained embedding space. Such a model would also grow in complexity as the inputs towards the neural networks begin to integrate multi-model data types. Advances in single cell manipulation has allowed for more platforms that allows for simultaneous measurements from within a single cell such as protein marker expression, chromatin occupancy, DNA mutations. We foresee the integration of such multi-model data types alongside with clinical phenotypes in a neural network-based prediction model to aid clinical management of the future.

## METHODS

### Siamese Network Architecture and Training

#### Architecture

The architecture of the Siamese network as its name implies has two inputs vectors **X**_**1**_, **X**_**2**_ that feeds into a common neural network that shares the same weights ***W***. This dense fully connected neural network consists of an input layer with 33694 nodes, each corresponding to a specific gene, followed by 2 fully connected layers each with 512 hidden nodes and a final 32 nodes output layer. The final output layer represents a 32-dimension feature space that is intended for separation of different cell types. A custom distance layer takes the transformed vectors and calculates the Euclidean distance in this embedded space:

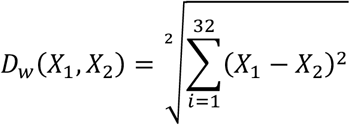

#### Generalizability

Between each fully connected layer, an additional dropout layer at a rate of 0.5 is implemented during training to ensure generalizability of the network during implementation. This network is implemented and trained using Keras and TensorFlow in both R and python environments.

#### Training with Contrastive Loss

35 cells are randomly selected from each cell type. Selected cells are split into training (20 cells per type) and validation sets (15 cells per type). Across the selected 20 cells of each types, pairs are generated: pairs originating from same cell type are labelled as 1 and pairs of cells from different cell types are labelled as 0. Gene counts of each cell are normalized by scaling with the maximum gene count of the cell. The binary cell labels **Y**, and Euclidean distance of the two-feature vector derived above **D**_**w**_ is fed into the contrastive loss function:

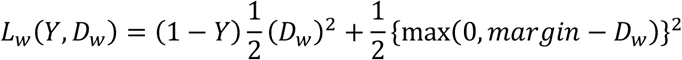

This loss was back-propagated to calculate the gradient and RMSprop ^14^ was used to update the weights.

### Visualization of training process

Visualization of the training process begins with calling back the weights of the neural networks across the training epochs. Weights corresponding to each training epoch are loaded, and each cell’s gene expression vector are passed through the network, where the final output of the embedding layer of a vector length 32 for each cell are collected and reduced into a two-dimensional space using UMAP. Individual firing of each of the 32 nodes in the final layer of neural networks are also visualized using heatmaps using the R package ComplexHeatmap^15^.

### Comparison of SNN distance with Euclidean & Cosine distance

20 cells are randomly selected from each of the annotated cell clusters of reference patient data. Each of these cells are paired with 20 other cells from the other annotated clusters. The distance between the 20 pairs of cells across the different annotated cell types are calculated using the SNN, Euclidean and cosine metric. The resulting distance for the distance metric is visualized using bar graphs in Figure 3. In order to quantitate the contrast in distance between the exact match and second-best match in terms of annotated cell types, we calculate the Signal to Noise Ratio between the top two matches:

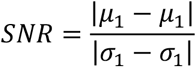

### Validation of SNN distance usage in annotating external datasets

Single cell gene expression data was obtained from the 10× genomics (https://support.10xgenomics.com/single-cell-gene-expression/datasets/3.0.2/5k_pbmc_v3). The dataset contains both gene expression and cell surface protein expression data from single cells. Each cell gene expression vector is matched up accordingly to the gene inputs that the Siamese model was trained on. Each of the external single cell gene vector is then paired against the trained reference cell types and fed into the Siamese network to obtain the SNN distance. The cell type of the reference cell group that correspond to the lowest SNN distance is then used to annotate the cell. Diagrammatic interpretation of the workflow is shown in figure 6.

### Validation of SNN distance usage in annotating different species

Single cell gene expression data from mouse samples in the same study was mapped to orthologous human genes using Mouse Genome Informatics MGI (http://www.informatics.jax.org/downloads/reports/HMD_HumanPhenotype.rpt). Human genes with no known mouse orthologs are set to zero. This transformed input is then paired against the trained reference cell types and fed into the Siamese network to obtain the SNN distance. The cell type of the reference cell group that correspond to the lowest SNN distance is then used to annotate the cell.

### Generalizability of Siamese trained embedding space in distinguishing cell types

Single cell gene expression vector from cell types not used during the training of the model were selected and embedded using the prior trained embedding neural net. K-nearest neighbor (KNN) is then performed on the feature vectors to generate a KNN graph network. Visualization of separations in the novel cell clusters within the network is achieved by using Fruchterman-Reingold force layout.

### Generation and Annotation using Human Cell Landscape atlas as a reference

Raw data was obtained from the human cell landscape portal [16]. Using the cell annotations provided, we tallied the different cell types within each tissue type. Only cell types, within each tissue, that have at least 30 cells were used for training. 20 cells are sampled from each cell type to generate pairs for training. The remaining 10 cells are used for validation. A binary indicator vector of same length to the number pairs is also generated where 1 indicates the pair of cells are drawn from the same cell type and 0 otherwise. The prepared data of cell pairs is fed into the SNN architecture as defined earlier.

For training the HCL dataset, the computational demand on hardware memory necessitated running the training on an AWS p2.large instance. All other training runs were performed on a local desktop with a RTX-2080 GPU. Callbacks were made to save the weights of the network at each epoch. To evaluate the progress of the training, a Siamese accuracy metric defined by arbitrarily setting the Euclidean distance at 0.5 where a distance lower than 0.5 is deemed that the cells are derived from the same cluster and conversely, distances greater than that are determined to be cells from different cell cluster. Weights from the epoch that gives the highest achievable training and validation accuracy are retained for deployment during annotation phase. Using the learned embedding from the network, the dimension reduced vectors of these reference cell groups are used to generate a reference KNN network. For the annotation phase, each of single cell vector from the Zilionis dataset is projected into the same space, and annotation is transferred using K nearest neighbor with K set at 3.

### Generation of Digital Twins via embedding space of SNN

Using the same process of training the human landscape model, the process is repeated across each of the patients in the Zilionis dataset. A different embedding space is derived from each of the patient’s trained network. Each of these embedding spaces is used to annotate the same held-out patient test dataset. Comparisons of the resulting cell type annotation from using the different embedding schemes are visualized using alluvial plots in R using ggalluvial package.

### Deployment of SNN networks as a restful API for community usage

Enabling and abstracting the ease of annotation using complex neural network models for the larger scientific community is a major goal of our work. Towards this aim, we have hosted the trained model on AWS and has provided an API [http://13.229.250.159:8000/_swagger_/] that abstracts away the need for complicated deployment for annotation. Each http GET request will send a JSON formatted single cell gene vector to the API which will annotate a single cell within 300 seconds, below the timeout limit (900 sec) of AWS lambda functions. While this cloud deployment scheme, will be slower in deployment compared to a local server model, we believe that a cloud deployment allows for much easier access to the trained model and has the scalability to better serve the wider community.

### Interspecies Annotation using SNN

To use the human trained SNN model for mouse annotation, we first obtained the mouse-human orthologs from Mouse Genome Informatics (MGI) [http://www.informatics.jax.org/downloads/reports/HMD_HumanPhenotype.rpt]. Single cell RNA-seq data from mouse with a human orthologs are mapped to the same input using the SNN. The rest of the human gene inputs with no corresponding mouse orthologs are set to zero. The resulting inputs are compared to the human reference cell with known annotations and the three nearest reference human cells in the embedded space identified by K nearest neighbor were used to annotate the mouse input cell.

## ACKNOWLEDGMENTS

We would like to acknowledge the late Dr. Sydney Brenner for his inputs, advise and inspiration for this work. We would like to acknowledge the larger single cell community for providing the wealth of data that makes such an computational exploration possible.

